## Supplementary figures and images for "A phosphate-binding pocket in cyclin B3 is essential for XErp1/ Emi2 degradation in meiosis I"

### Supplemental Figures

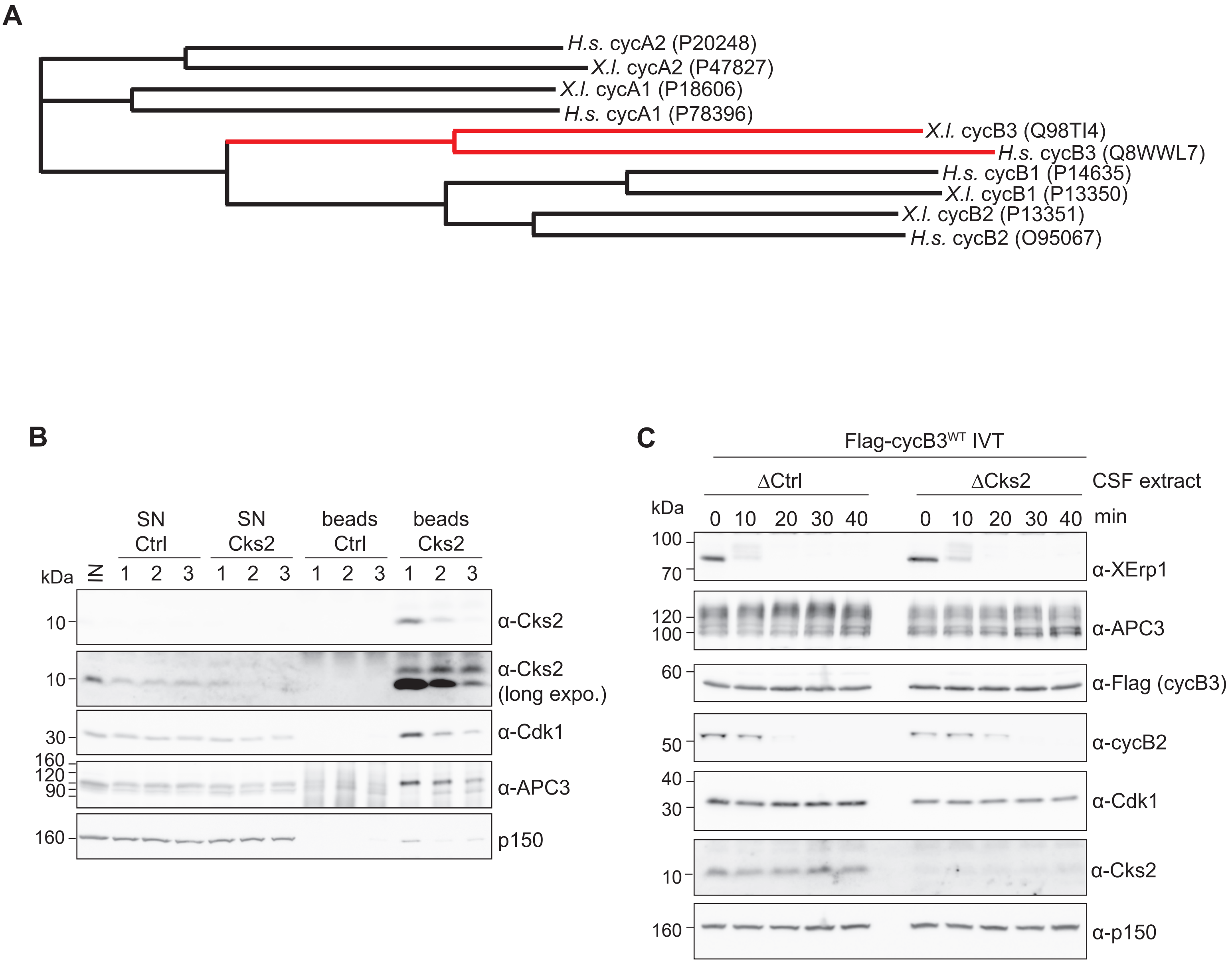

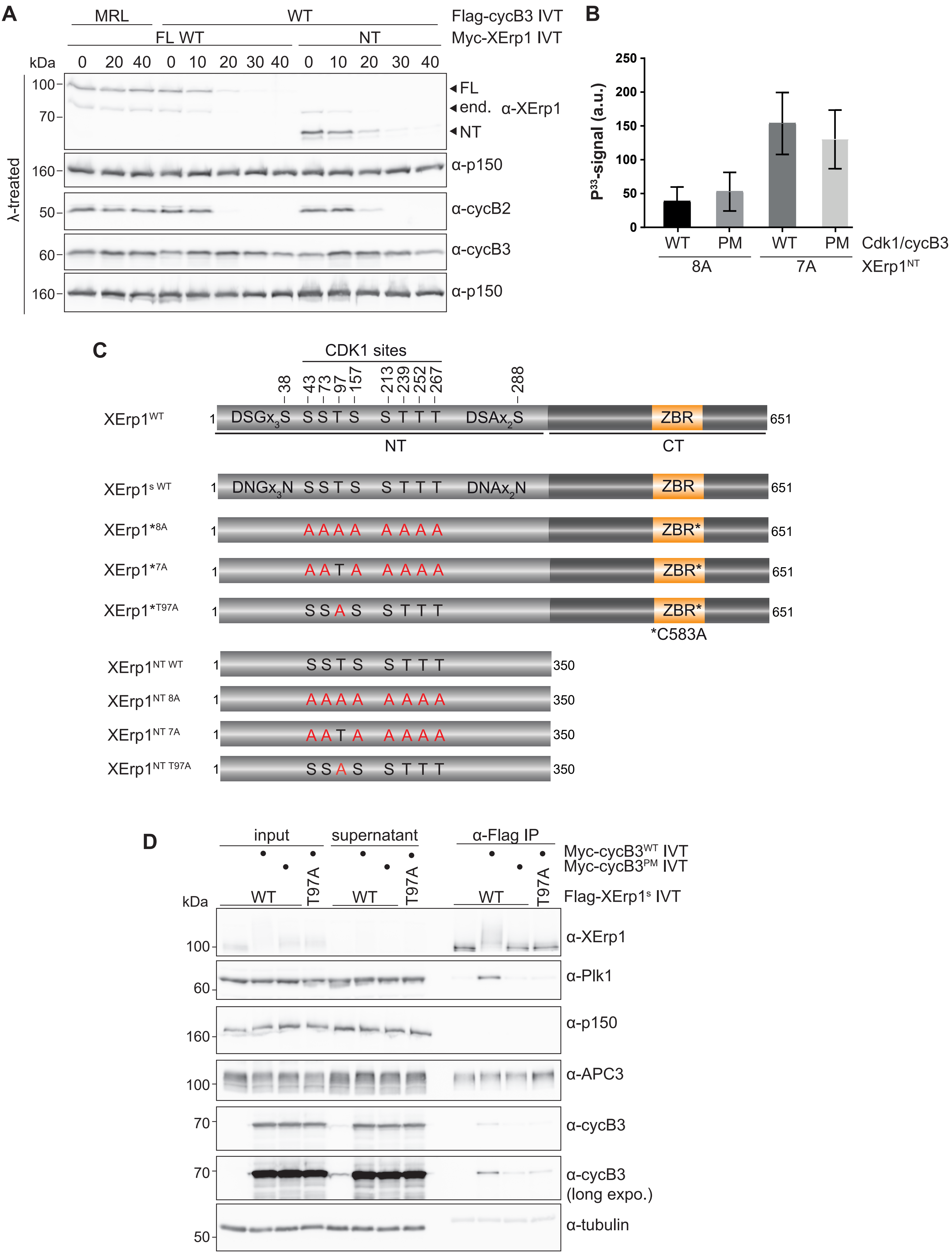

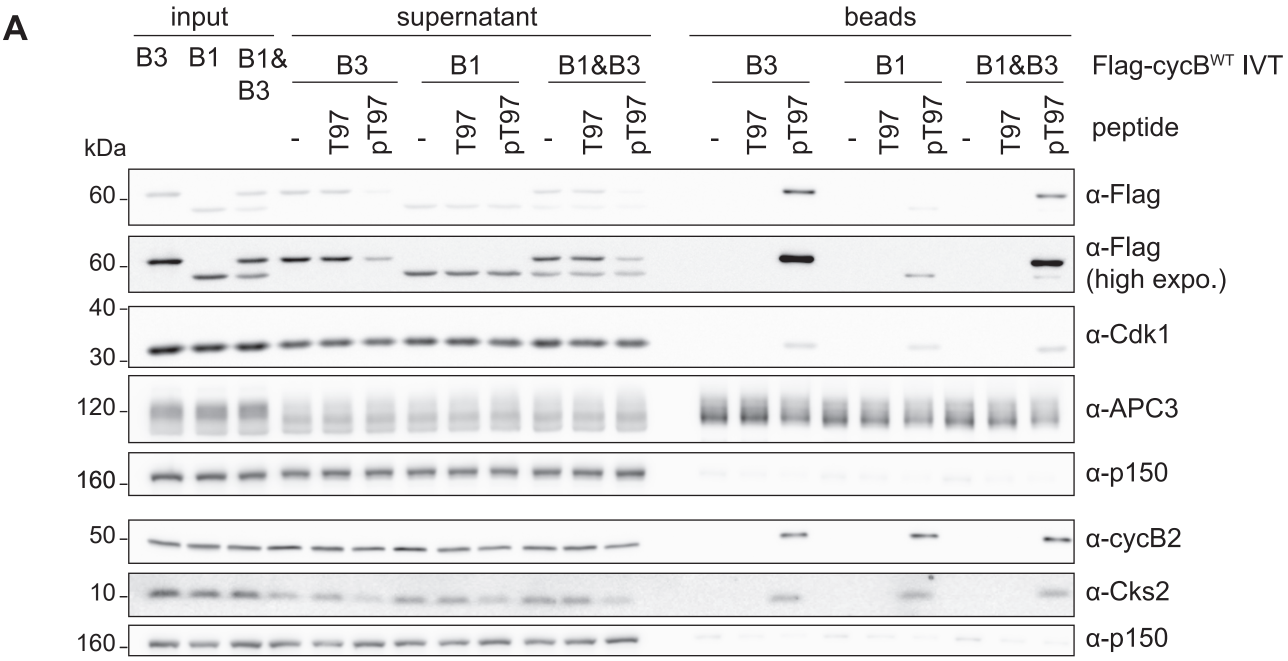

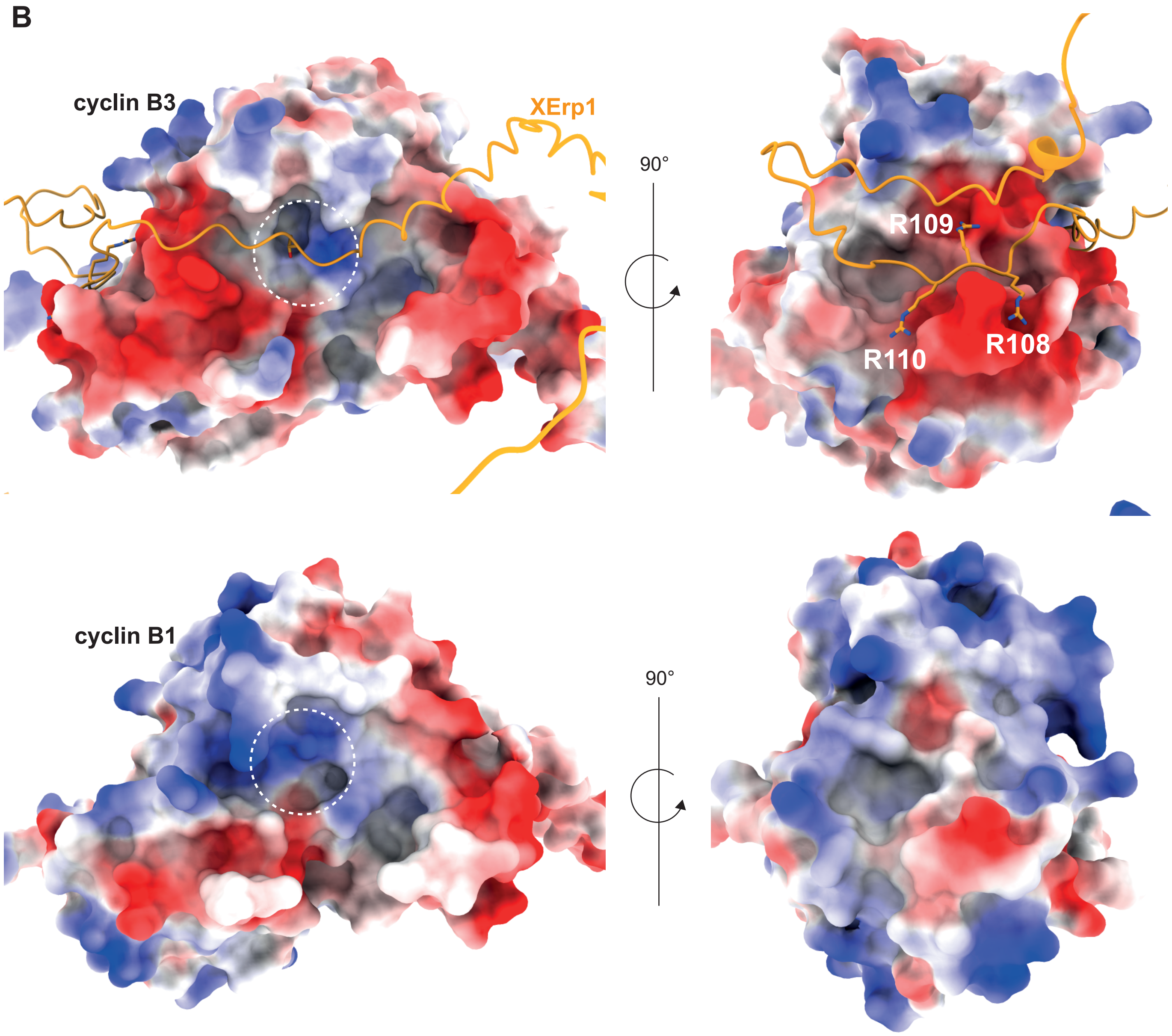
